# Constraining activity and growth substrate of fungal decomposers via assimilation patterns of inorganic carbon and water into lipid biomarkers

**DOI:** 10.1101/2023.11.21.568133

**Authors:** Stanislav Jabinski, Wesley d.M. Rangel, Marek Kopáček, Veronika Jílková, Jan Jansa, Travis B. Meador

## Abstract

Fungi are among the few organisms on the planet that can metabolize recalcitrant C but are also known to access recently produced plant photosynthate. Therefore, improved quantification of growth and substrate utilization by different fungal ecotypes will help to define the rates and controls of fungal production, the cycling of soil organic matter, and thus the C storage and CO_2_ buffering capacity soil ecosystems. This study combined a dual stable isotope probing approach together with rapid analysis by tandem pyrolysis gas chromatography isotope ratio mass spectrometry (Pyr-GC-IRMS) to determine the patterns of water-derived hydrogen (H) and inorganic C assimilated into lipid biomarkers of heterotrophic fungi as a function of C substrate. The water H assimilation factor (α_w_) and the inorganic C assimilation for C_18:2_ fatty acid isolated from five fungal species growing on glucose was lower (0.62±0.01 and 4.7±1.6%, respectively) than for species grown on glutamic acid (0.90±0.02 and 7.4±3.7%, respectively). Furthermore, the assimilation ratio (R_IC/αW_) for growth on glucose and glutamic acid can distinguish between these two metabolic modes. This dual SIP assay thus delivers estimates of fungal activity and may help to delineate the predominant substrates that are respired among a matrix of compounds found in natural environments. (200 Words)

**Importance:** Fungi decomposers play important roles in food webs and nutrient cycling because they can feed on both labile and stable forms of carbon. This study developed and applied a dual stable isotope assay to improve the investigation of fungal activity in the environment. By determining the incorporation patterns of hydrogen and carbon into fungal lipids, this assay delivers estimates of fungal activity and the different metabolic pathways that they employ in ecological and environmental systems. (75 Words)

## 1. Introduction

The relatively large fluxes between atmospheric and terrestrial carbon (C) reservoirs determine climate-soil feedbacks, especially because soils are highly sensitive to global and local climate variability^1^. As such, small changes in soil C reservoirs may have big impacts on atmospheric CO_2_ concentration^2–4^ and are one of the largest unknowns in the global C budget^5^. Uncertainty in future climate models can be improved with increased understanding of the activity of organisms that decompose the wide variety of substrates that constitute soil organic matter (SOM)^6–8^. While most studies of SOM dynamics have focused on bacteria, fungi are known to be one of the major decomposers in soils and may play a unique but underappreciated role^9^. Fungi are known to feed on chemically stable forms of organic C, such as cellulose^10^ and lignin^11–13^, and are also known to directly access recently produced plant photosynthate^14,15^.

Heterotrophic microorganisms who mainly feed on organic substrates also fix a variable amount of inorganic carbon. This flux acts to replenish intermediates in the tricarboxylic acid (TCA) cycle that have been released for biosynthesis^16^. It has been suggested that 2 - 8% of heterotrophic biomass originates from CO_2_ incorporated via carboxylation reactions^17–19^, largely depending on the redox state of the organic substrate. Although these processes were described decades ago^16,20^ the relevance and the (metabolic) controls on how much inorganic carbon is fixed by heterotrophic microorganisms is poorly understood, due to the lack of reliable estimates for most organisms and habitats^19^. Despite their prevalence as decomposers, estimates of inorganic carbon fixation by heterotrophic fungi are scarce^20^.

In recent years, hydrogen (H) isotope incorporation from water was shown to be a useful tracer of microbial community activity in a diverse range of environments^21–23^, as water is required by all microbes and there exists a strong relationship between the stable H isotopic composition of microbial lipids and water, which is the ultimate source of H in all-natural organic compounds^24,22^. Strong correlations between H isotopes of water medium and microbial lipids were shown to correlate with the type of growth metabolism^25^, with slightly higher δ^2^H values for aerobic heterotrophs, to lower δ^2^H values for phototrophs, to lowest δ^2^H values for chemoautotrophs^25^.

To fully exploit the potential of stable isotope probing (SIP) experiments, a new lipid-based dual-SIP approach was developed to track total microbial production by adding heavy water (D_2_O) together with labeled ^13^C-dissolved inorganic carbon (DIC), to enable estimates of total and autotrophic metabolism, respectively^26,27^. Because bicarbonate is often present in relatively high concentrations in natural systems, adding ^13^C-bicarbonate and D_2_O to incubation experiments does not alter ambient biogeochemical conditions^26^. The ratio of D_2_O incorporation and ^13^C-DIC assimilation into lipid biomarkers can then distinguish microbial metabolism of autotrophs (ratio ∼ 1) and heterotrophs (ratio < 0.3)^26^.

The purpose of this work was to apply the dual stable isotope approach to determine heterotrophic IC fixation and the water H assimilation factor for fungal lipid biomarkers produced during growth on labile substrates (glucose and glutamic acid), with the hypothesis that growth substrate will activate alternative metabolic pathways with characteristic assimilation ratios of ^13^C versus ^2^H. We further tested the applicability of pyrolysis coupled gas chromatography – mass spectrometry (GC-MS) and isotope-ratio mass spectrometry (IRMS) as a rapid, direct analysis of isotope incorporation into fungal biomarkers.

## 2. Results

All five microscopic fungal species *Aspergillus dimorphicus* (AD), *Cordyceps farinosa* (CF), *Penicillium janczewskii* (PJ), *Acremonium polycromum* (AP) and *Aspergillus brasiliensis* (AB) and the yeast species *Yarrowia lipolytica* (YL) exhibited a capacity to grow in the modified minimum media Czapek-Dox Broth (Thom & Raper 1945), with either glucose or glutamic acid as a sole carbon source, and having various enrichment levels of ^13^C-DIC and D_2_O.

### 2.1 Fungal growth

CO_2_ concentrations in the fungal incubations increased linearly and the biomass was harvested when the CO_2_ headspace concentration reached 4 – 6% v/v (Fig. 1), which required incubations times ranging 2 – 4 weeks for glucose and 4 – 8 weeks for glutamic acid. Incubations with CF grown on glucose were performed in 1 L bottles and reached higher CO_2_ concentrations of ∼ 10%. YL grown on glucose did not show any increase in CO_2_ during the incubation period of 6 weeks. d^13^C of headspace CO_2_ decreased from a maximum value of +5000‰, measured 3 days after inoculation, to a minimum of approximately +40‰ at the end of the incubation, as CO_2_ released during the respiration of unlabeled glucose or glutamic acid slowly diluted the ^13^C-label. Biomass harvested from each incubation bottle amounted to 50-100 mg dry weight.

**Figure 1:**
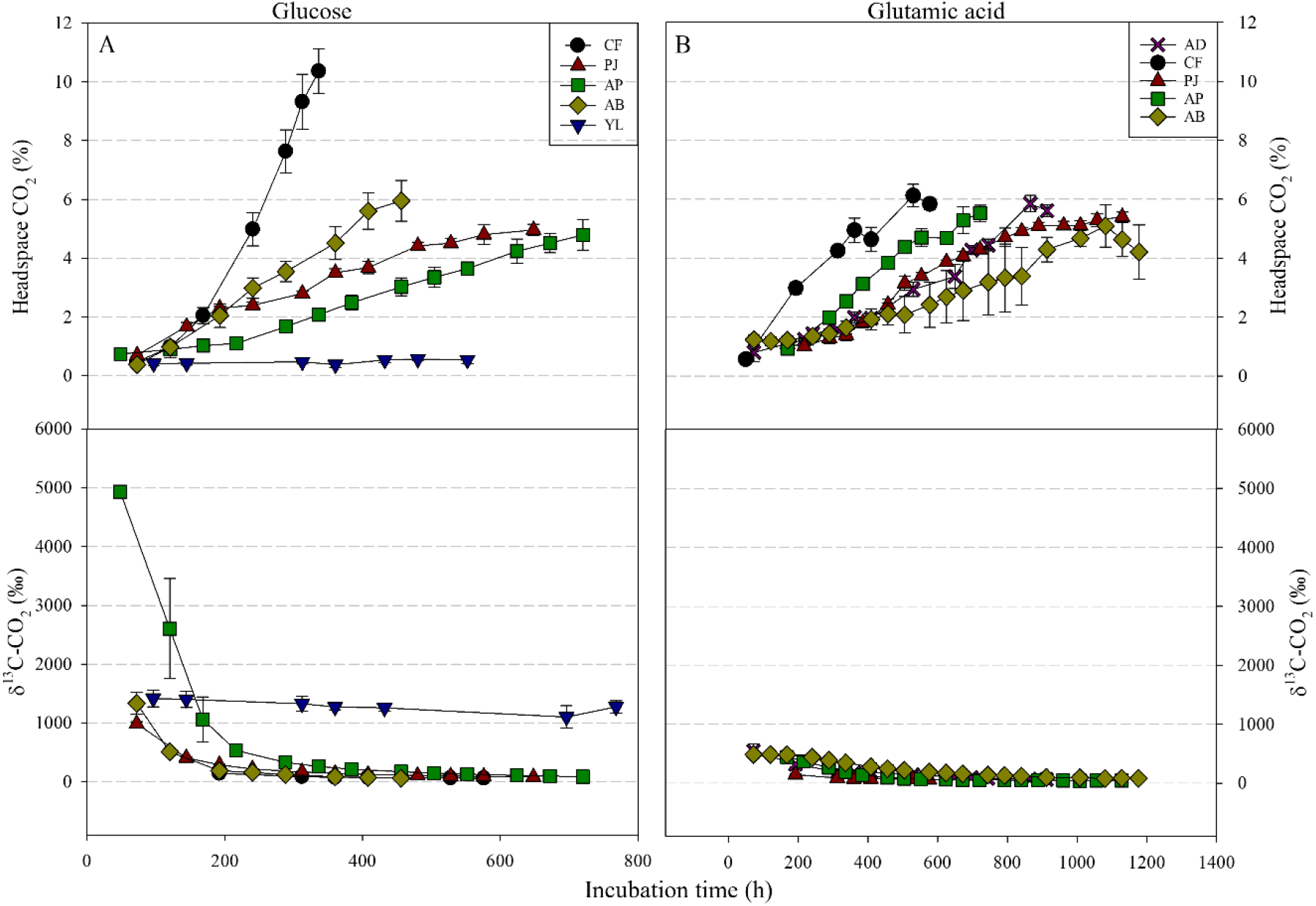
Headspace CO_2_ concentrations for the fungal incubation grown on glucose (A) and glutamic acid (B) over time with their respective δ^13^C-CO_2_ values in the lower panels. Incubation was stopped when CO_2_ concentrations in the headspace reached 4 - 6% (v/v), which is indicative that roughly half of the carbon substrate was respired, and based on a mol-to-mol comparison, ensures the incubation was still under aerobic conditions. AP = *Acremonium polycromum*, AB = *Aspergillus brasiliensis*, AD = *Aspergillus dimorphicus*, CF = *Cordyceps farinosa*, PJ = *Penicillium janczewskii* and YL = *Yarrowia lipolytica*.

### 2.2 Analysis of extracted fungal lipids by GC-FID/MS/IRMS

#### 2.2.1 Fatty acid distributions

The fatty acid composition of all strains were similar, except for YL, which contained only four different fatty acids (Fig. 2). The fatty acids C_16:0_, C_18:0_, C_18:1_ and C_18:2_ accounted for about 80 – 100% of the total lipid content when grown on glucose. The fatty acids C_16:0_ and C_18:2_ accounted for 80 – 94% of the total lipid content when grown on glutamic acid. Higher fatty acid diversity was found for fungi grown on glucose *versus* those grown on glutamic acid.

**Figure 2:**
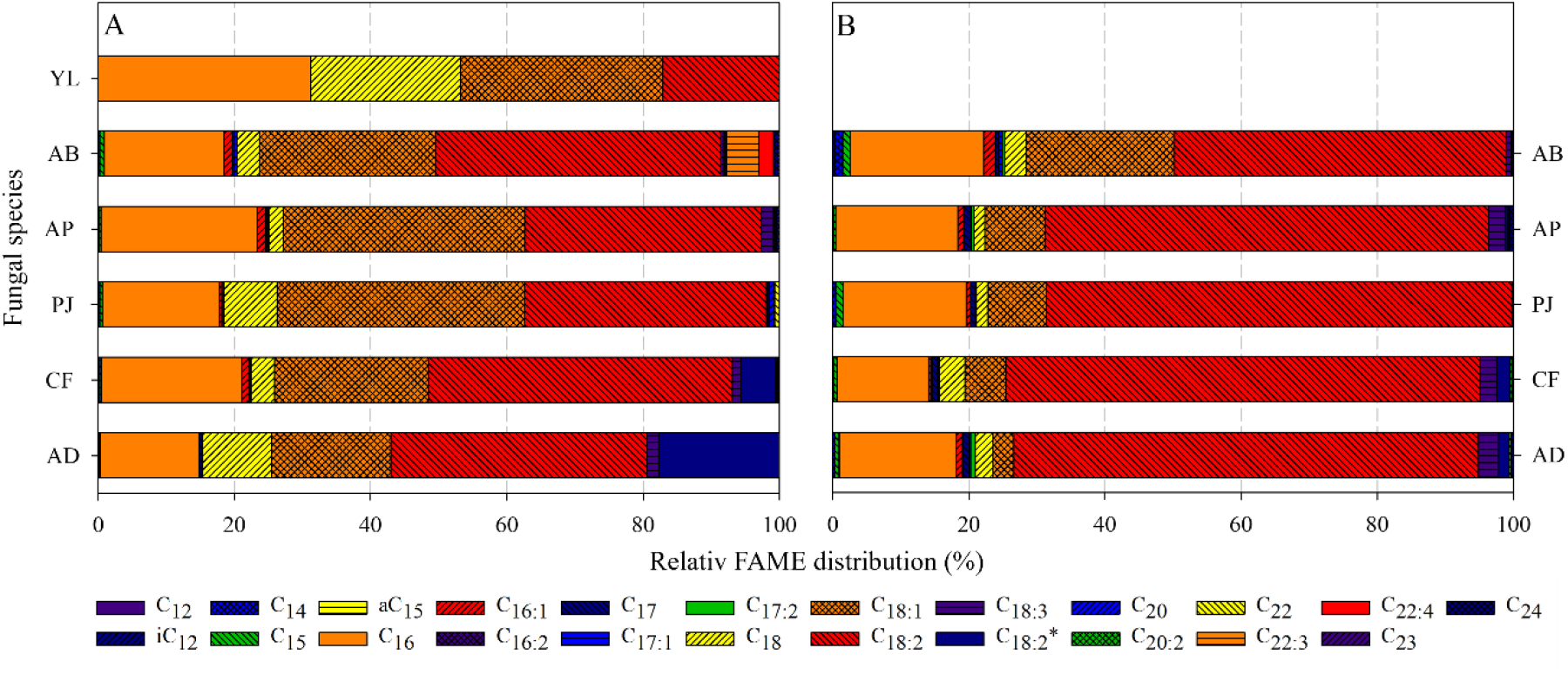
The fatty acid composition for all strains grown on glucose (A) and glutamic acid (B), and determined by conventional GC-FID. A slightly higher diversity of fatty acids was found when grown on glucose with the dominant fatty acids C_16:0_, C_18:0_, C_18:1_ and C_18:2_ accounting for more than 80%. The dominant fatty acids when grown on glutamic acid were C_16:0_ and C_18:2_, accounting for more than 80% of total fatty acid content. C_18:2_ represents a fatty acid with two unsaturations in the C_18_ chain at positions 9 and 12, whereas C_18:2_* represents the sum of all other isomers with two unsaturations in the C_18_ chain. AP = *Acremonium polycromum*, AB = *Aspergillus brasiliensis*, AD = *Aspergillus dimorphicus*, CF = *Cordyceps farinosa*, PJ = *Penicillium janczewskii* and YL = *Yarrowia lipolytica*.

#### 2.2.2 Carbon isotopes

The δ^13^C values of fungal biomarkers C_18:2_ and ergosterol produced under natural cultivation conditions with glucose (i.e., non-labeled; AT%_DIC_ ∼1%) ranged from −31.2‰ to −27.5‰ and - 28.6‰ to −24.8‰, respectively, across all strains. Fungal lipids were typically depleted in ^13^C relative to glucose (δ^13^C_glucose_ = −26.5‰), with an isotope effect [ε_lipids/glucose_ = ((δ_lipid_ + 1) / (δ_glucose_ +1) – 1) x 1000); cf.^28^] of −3.3±1.3‰ for C_18:2_ and −0.6±1.4‰ for ergosterol. The isotopic composition of C_18:2_ and ergosterol from the labeled incubation (AT%_DIC_ = 10%), displayed slightly higher δ^13^C values, ranging from −29.2‰ to −23.3‰ and −25.1‰ to −19.9‰, respectively. The apparent isotope effect of lipid biosynthesis therefore increased, yielding average ε_lipids/glucose_ values of +0.3±2.2‰ for C_18:2_ and +3.8±2.2‰ for ergosterol, roughly 4‰ higher than the non-labeled treatment.

The δ^13^C values of fungal biomarkers C_18:2_ and ergosterol produced under natural cultivation conditions with glutamic acid ranged from −20.8‰ to −17‰ and −18.4‰ to −13.9‰, respectively. Lipids were consistently depleted in ^13^C relative to glutamic acid (δ^13^C_glutamic acid_ = −12.7‰), with an isotope effect (ε_lipids/glutamic acid_) of −6.2±1.9‰ for C_18:2_ and −3.9±1.7‰ for ergosterol. The δ^13^C values of C_18:2_ and ergosterol from the labeled incubations (AT%_DIC_ = 10%) were higher, ranging from −14.8‰ to −10.3‰ and −14.9‰ to −9‰, respectively, and yielding ε_lipids/glutamic acid_ values (+0.7±2.5‰ for C_18:2_ and +1.7±3.3‰ for ergosterol) that were roughly 6‰ higher than non-labeled incubations.

The IC incorporation into fungal lipids (Fig. 3) was calculated after equation (Eq. 1).

**Figure 3:**
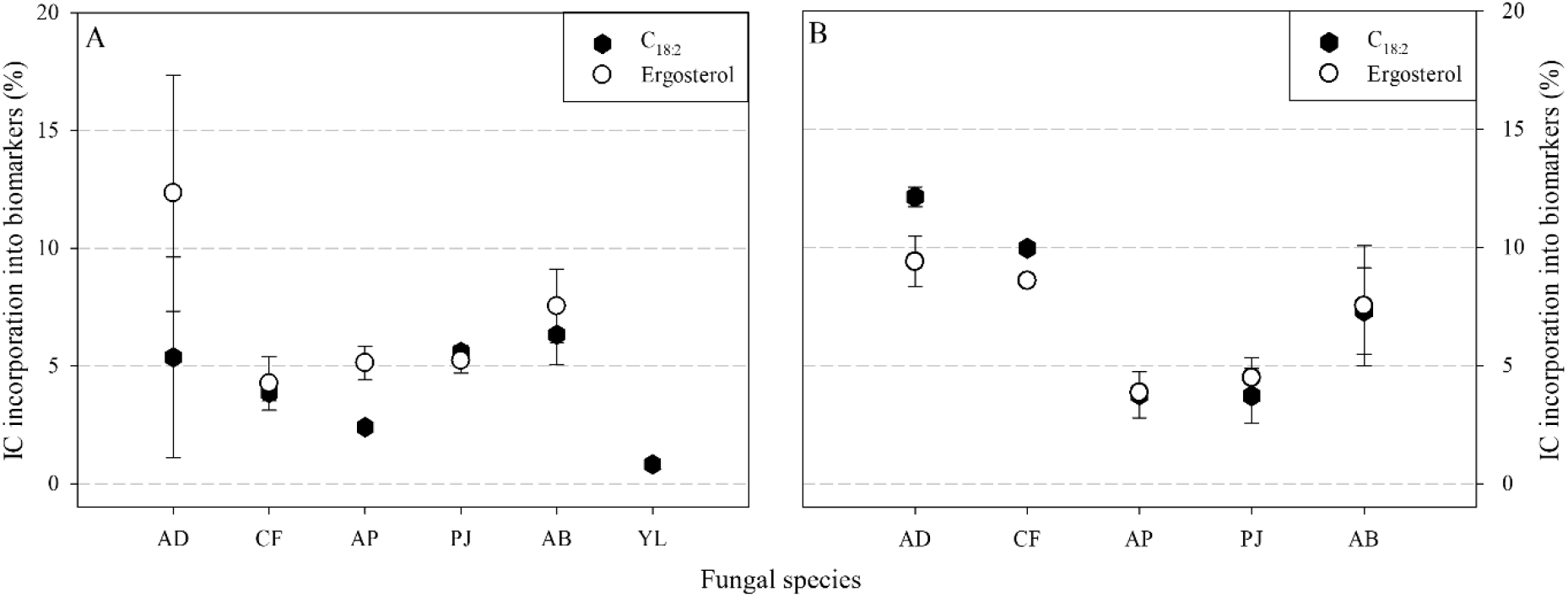
IC assimilation (y-axis) by fungal species (x-axis) into the specific biomarker fatty acid C_18:2_ and ergosterol incubated with glucose (A) and glutamic acid (B). Data are shown only for conventional GC-IRMS analyses following lipid extraction and purification (Section 5.2). AP = *Acremonium polycromum*, AB = *Aspergillus brasiliensis*, AD = *Aspergillus dimorphicus*, CF = *Cordyceps farinosa*, PJ = *Penicillium janczewskii* and YL = *Yarrowia lipolytica*.

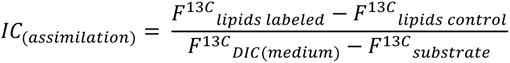

Equation 1: Inorganic carbon (IC) assimilation was calculated as the difference in the fraction of ^13^C [F^13C^ = (^13^C)/(^13^C+^12^C)] of the lipids harvested from the labeling experiment compared to the natural (control), normalized by the difference in F^13C^ of the DIC measured at the end of the incubation and the F^13C^ of the substrate. F was calculated as F^13^C = (R^13C/12C^)/(R^13C/12C^ + 1), where R is calculated from the δ^13^C ratios as measured with the IRMS equipment using the reverse of the δ notations (δ^13^C = ([^13^C/^12^C]_sample_/[^13^C/^12^C]_ref_ - 1) * 1000 (modified after ^32,26^).

For fungi growing on glucose, the average incorporation of IC into C_18:2_ and ergosterol was 4.7±1.6% and 6.9±3.3% respectively. When growing on glutamic acid the average IC incorporation into C_18:2_ and ergosterol was 7.4±3.7% and 6.8±2.5%.

#### 2.2.3 Hydrogen isotopes

The contribution of water-hydrogen to lipid biosynthesis was estimated in triplicate for fungi growing in natural MilliQ (−66±1‰) and three different enrichment treatments (+107±21‰, +209±18‰ and +411±11‰). A linear regression was then used to constrain the fractional abundance of ^2^H (F^2H^) between the medium water and the lipids, yielding a slope that represents the “net” contribution of water-derived H to lipid H, known as the water hydrogen assimilation factor α_W_ (Fig. 5;^25,22^). The α_W_ values for fungal biomarkers C_18:2_ and ergosterol harvested from glucose incubations ranged from 0.58±0.03 to 0.67±0.03 and 0.46±0.03 to 0.75±0.03, respectively. C_18:2_ harvested from the yeast strain (YL) growing on glucose exhibited a lower α_W_ value of 0.32±0.03 (Fig. 4); these data are excluded from subsequent calculations. The α_W_ values for fungal biomarker C_18:2_ and ergosterol from glutamic acid incubations ranged from 0.75±0.02 to 1.15±0.10 and 0.54±0.02 to 0.74±0.02, respectively. The average α_W_ value for C_18:2_ was significantly higher with glutamic acid as the C substrate (0.90±0.02; n = 58) *vs.* glucose (0.62±0.01; n = 42 ; *p* < 0.005), whereas the average α_W_ values for ergosterol were very similar between glutamic acid (0.68±0.08) and glucose (0.64±0.13).

**Figure 4:**
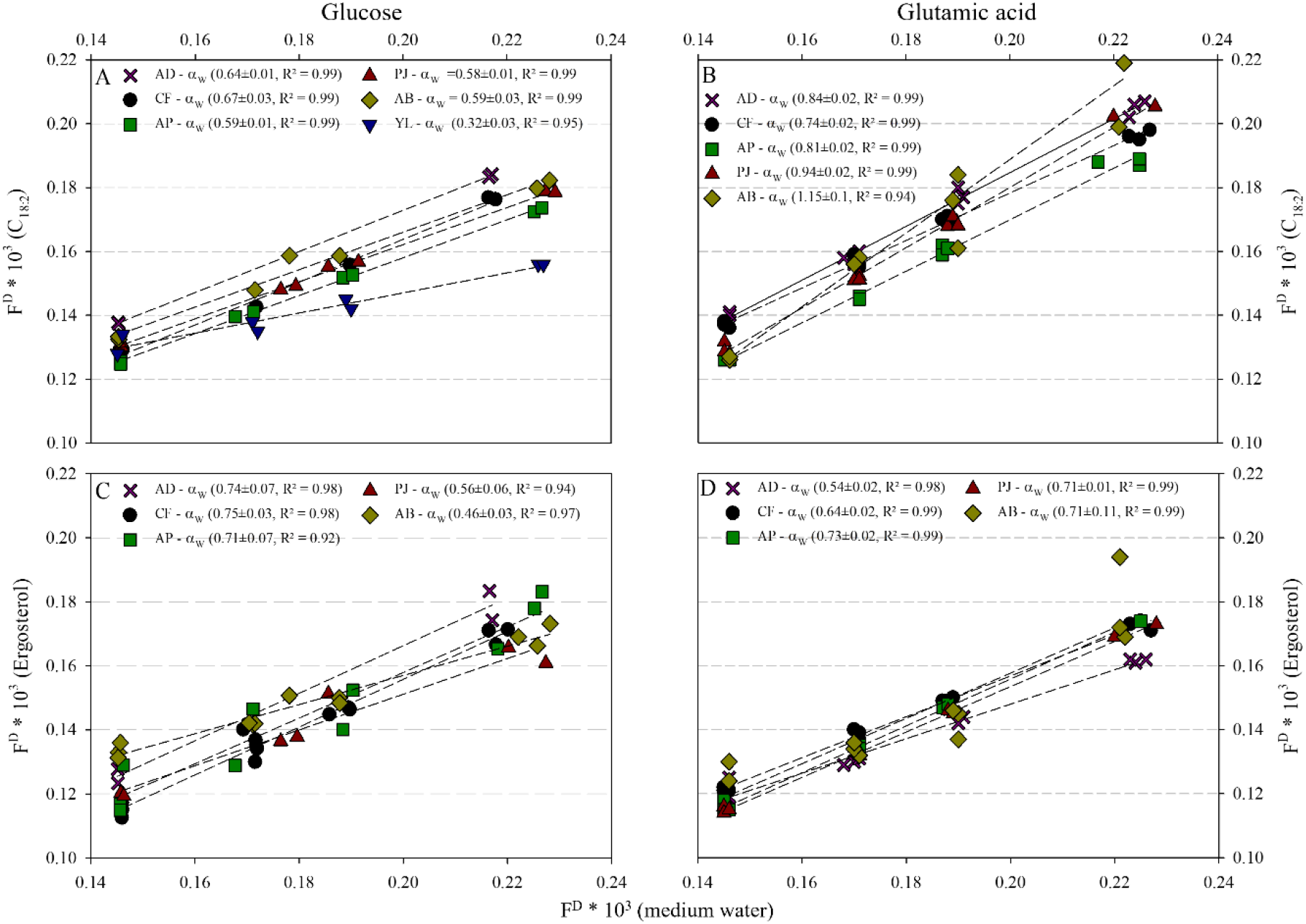
The water hydrogen assimilation factor (α_W_ values) estimated as the slope of the fractional ^2/1^H abundance (F^D^ × 10^3^) in lipids (y-axis) versus medium water (x-axis). Data are shown for fungal biomarkers C_18:2_ (top panels A and B) and ergosterol (bottom panels C and D) extracted from the different fungal species grown on glucose (A, C) or glutamic acid (B, D) using the conventional GC-IRMS method (Section 5.2). AP = *Acremonium polycromum*, AB = *Aspergillus brasiliensis*, AD = *Aspergillus dimorphicus*, CF = *Cordyceps farinosa*, PJ = *Penicillium janczewskii* and YL = *Yarrowia lipolytica*.

**Figure 5:**
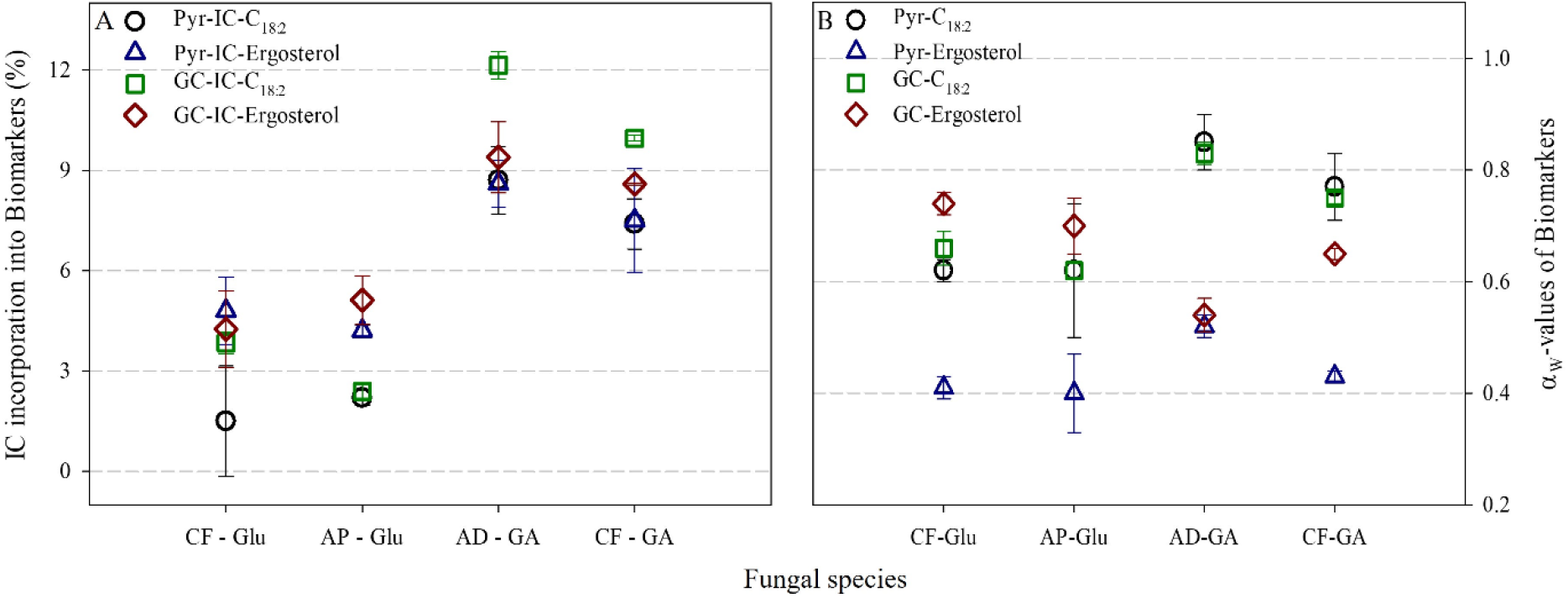
Comparison between Pyrolysis- and conventional GC-IRMS analysis of fungal biomarkers to determine IC incorporation (panel A) and water assimilation factor (α_W_) (panel B). AP = *Acremonium polycromum*, AD = *Aspergillus dimorphicus*, CF = *Cordyceps farinosa*.

### 2.3 Analysis of fungal biomass by Pyr-GC-FID/MS/IRMS

Four dual SIP experiments were selected to compare IC incorporation and α_W_ values determined via the extraction-free, pyrolysis approach (Pyr-GC-IRMS; n = 24 analyses) to the well-established, wet chemistry approach (GC-IRMS; n = 48 analyses). This included the species CF and AP grown on glucose and AD and CF grown on glutamic acid (Fig. 5; S-Tab. 2). The IC incorporation into the fungal biomarkers C_18:2_ and ergosterol from the glucose experiments were on average 3.1±1% and 4.6±0.6% (GC) *vs*. 1.9±0.5% and 4.5±0.4% (Pyr). For the glutamic acid experiment, the average IC incorporation into C_18:2_ and ergosterol were 11.1±1.5% and 9.0±0.6% (GC) *vs*. 8.1±0.9% and 8.1±0.8% (Pyr), respectively. The α_W_ values for C_18:2_ and ergosterol from the glucose experiment were on average 0.63±0.05 and 0.73±0.02 (GC) *vs*. 0.62±0.01 and 0.39±0.03 (Pyr). For the glutamic acid experiment, the average α_W_ values for C_18:2_ and ergosterol were 0.79±0.05 and 59.0±0.07 (GC) *vs*. 0.81±0.06 and 0.48±0.06 (Pyr).

## 3. Discussion

We have generated complementary datasets of IC and water H incorporation into fungal lipids using two different approaches. The interpretation of these estimates in the sections below are based on data determined via classical GC analysis, following extraction, purification, and transesterification of lipids as described in Section 5.2, and thus comparable with previous studies that have used this approach (e.g.,^25,22,33^). We further compare these values with those derived from direct analysis of fungal biomass via pyrolysis-GC (cf. Section 5.3.5), as this approach may promote higher throughput for lipid analyses and thereby help to constrain IC and water H assimilation factors across more species and carbon substrates.

### 3.1 IC assimilation into fungal lipids

CO_2_ assimilation by heterotrophic organisms is reported to be a measure of both anabolic processes and the catabolic status of the cell, with assimilation, anaplerotic, biosynthesis, and redox balancing reactions playing important roles to ensure the provision of energy and to replenish intermediates in the tricarboxylic acid (TCA) cycle that have been released for biosynthesis^16,34,35^. The by-fixation of inorganic carbon via anaplerotic pathways typically varies between 1-8% of the carbon biomass, depending on central metabolic pathways, and could further increase to assimilate carbon substrates that are more reduced than the biomass, as various carboxylase enzymes are employed to match the redox state of the substrate with that of the biomass^16^. Whereas IC assimilation by fungi was previously reported to amount to roughly 1%, based primarily on radiocarbon uptake into whole cells^20,36,37^, our results suggest that ^13^C-labelled IC accounted for 2 – 12% of C assimilated into fungal membrane lipid biomarkers, when grown on minimum medium with glucose or glutamic acid as the main carbon source. Both glucose and glutamic acid are slightly more reduced (degree of reduction, DOR = 4 and 3.6, respectively^38^) than expected for biomass (DOR = 4.25;^19^), suggesting that assimilatory carboxylation should not contribute as strongly as anaplerotic or biosynthesis reactions to IC by-fixation during growth on theses substrates. Our findings indicate that ^13^IC incorporation into the C_18:2_ biomarker was slightly lower during growth on glucose substrate (4.7±1.6%) versus glutamic acid (7.4±3.7%; Fig. 3), which may have involved more carboxylation reactions to replenish central metabolites. For example, during growth on glutamic acid, relatively more carboxylase reactions may be required to replenish acetyl-CoA, thereby increasing the conduit for by-fixation of IC. Consistent with this notion, growth on glutamic acid was slower than glucose across fungal species (Fig. 1).

In contrast to the observations for C_18:2_, ^13^IC incorporation into ergosterol was similar for both substrates (6.9±3.3% and 6.8±2.5%; Fig. 3). The different patterns of IC incorporation into C_18:2_ versus ergosterol likely owes to their different biosynthetic pathways, where both products start from acetyl-CoA. For fatty acids, acetyl-CoA molecules are merged to malonyl-CoA, and other acetyl-CoA molecules are added repeatedly until the desired chain length for the fatty acid is attained. Ergosterol is synthesized from squalene (C_30_), which is produced by merging C_5_ intermediates of the mevalonate pathway (e.g., isopentenyl-pyrophosphate); each C_5_ unit of squalene stems from the merger of three acetyl-CoA molecules a decarboxylation reaction. Squalene further undergoes cyclization to form lanosterol, the main precursor for sterols, through the subsequent removal of three methyl groups^39,40^. Before the final ergosterol molecule is produced, another methyl group is added to the backbone^39,40^. The removal and/or replacement of acetyl-CoA-derived C during ergosterol biosynthesis may thus make this biomarker less-sensitive than fatty acids to the ^13^IC incorporation via assimilation or anaplerotic pathways.

### 3.2 Water hydrogen incorporation into fungal lipids

Previous studies have demonstrated that the regression slope between hydrogen isotopes of water medium and microbial lipids (i.e., α_W_) varies with the type of metabolism used during growth and different organisms may exhibit similar fractionation or α_W_ values when grown on the same substrate^25,41,42^. Variability in fatty acid hydrogen isotopic composition is suggested to be a function of the isotope effects of hydrogen transporters and electron acceptors (NADPH and NADH), as they are the main portals for hydrogen incorporation in FA biosynthesis pathways (accounting for around 50%), with the remaining coming directly from environmental water (∼ 25%) and acetyl-CoA (∼ 25%)^25,42^. In the current study, the water-derived hydrogen contribution to membrane lipids was consistent among fungal species grown on the same substrate and similar to the patterns observed for IC incorporation, with the average α_W_ value of the C_18:2_ fatty acid biomarker harvested from glucose incubations being significantly lower than glutamic acid incubations (0.62±0.04 versus 0.90±0.16, *p* < 0.005). This finding indicates that the central metabolism of glutamic acid resulted in a higher contribution of H from environmental water to C_18:2_. For ergosterol, α_W_ values for glucose and glutamic acid incubations were not significantly different (0.64±0.13 and 0.68±0.08, respectively), suggesting that ergosterol is less sensitive to H contributions from water in the growth medium. In the context of previously reported α_W_ values for the three domains of life, fungal lipids are more comparable to that of bacteria bacteria (0.4 – 0.9) than that of eukaryotes (0.7 – 1.2), although only phyto- and zooplankton α_W_ values are reported so far for that domain (^43^ and references within).

### 3.3 Dual-SIP approach

Dual SIP experiments with HDO and ^13^C-DIC have been applied to distinguish autotrophic versus heterotrophic modes of carbon assimilation for pure cultures of bacteria and archaea^26,27^ and were also used to track microbial activity in marine sediments^21,44,45,33^. The main objective of this study was to develop the dual-SIP assay to determine ^13^IC and ^2^H incorporation into fungal lipid biomarkers (Ratio ^13^IC/α_W_), which could be further applied to more comprehensively explore the patterns derived from different functional and/or taxonomic groups of fungi. The average ^13^IC/α_W_ ratios for C_18:2_ were 0.076±0.026 and 0.11±0.057 for glucose and glutamic acid, respectively. This first evidence defines distinct spaces for fungal growth on the different substrates in the ratio plot (Fig. 6) and supports the use of the dual SIP assay to distinguish the different metabolic modes employed by fungi for growth on sugars versus amino acids. For ergosterol, the ^13^IC and α_W_ values determined for both carbon sources could not be distinguished in the ratio plot (Fig. 6), suggesting poor potential for ergosterol to serve as a biomarker of metabolic mode. More data and studies of additional taxa and C substrates are needed to fully exploit the potential of this approach.

**Figure 6:**
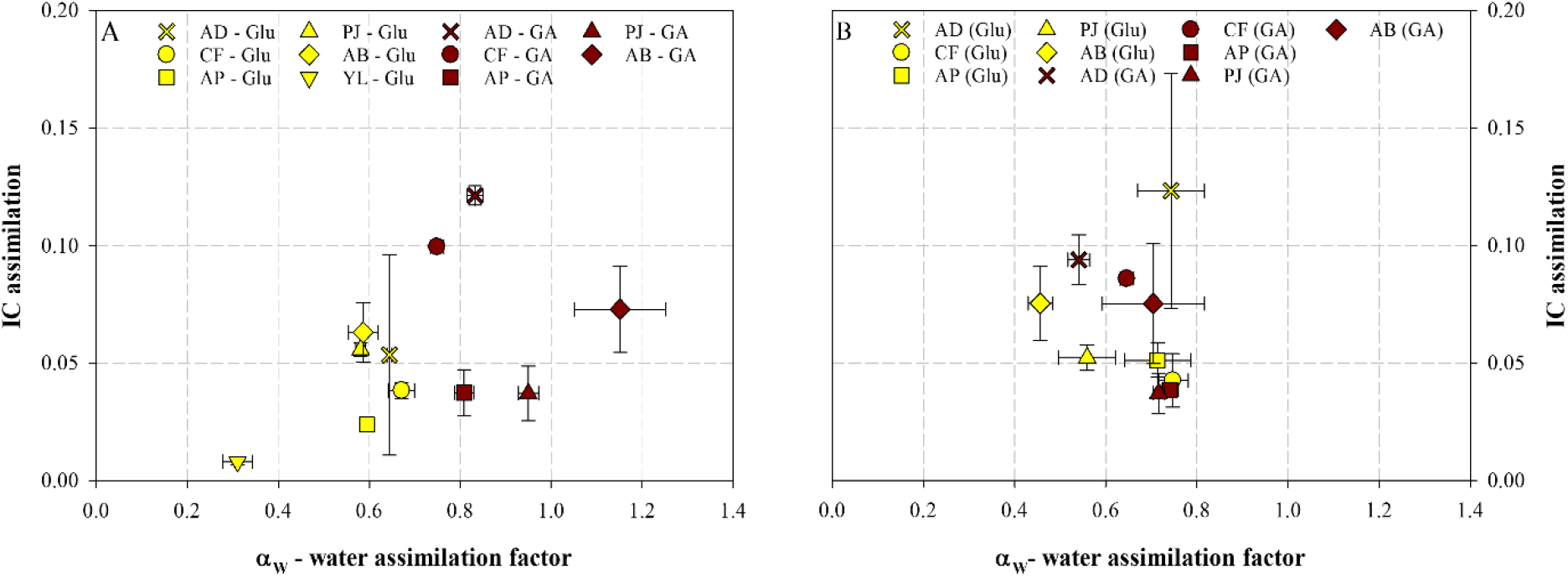
IC and α_W_-values ratio plots for C_18:2_ (A) and ergosterol (B). The power of the dual-SIP approach for fungal biomarkers shows distinguished groupings for glucose (yellow) vs glutamic acid (red) for C_18:2_, whereas these groups overlap for ergosterol.

### 3.4 Pyr-GC-MS/IRMS

An additional objective of this study was to test the potential of using Pyr-GC-IRMS to determine the isotopic composition of the fungal biomarker C_18:2_ and ergosterol for a rapid analysis that avoids time and solvent consuming wet chemistry steps. The application of Pyr-GC-MS for the analysis of whole cells was previously reported for fatty acid quantification^46^, which relies on thermal-assisted hydrolysis and methylation via a derivatization agent to enhance detection. In the current study, we used trimethylsulfonium hydroxide (TMSH) as a derivatization agent because it was reported to not cause any isomerization and/or degradation as observed for other TMH solutions^47,48^.

For the dual SIP experiments selected for both analyses, the patterns of ^13^IC incorporation and α_W_ values determined by the conventional GC-IRMS protocol were generally redundant with results from Pyr-GC-IRMS analysis, with the exception of α_W_ values calculated for ergosterol (Fig. 5; Table S2). The average IC incorporation determined for ergosterol were consistent between traditional GC-IRMS and Pyr-GC-IRMS protocols, and while the incorporation trends were consistent, ^13^IC incorporation into C_18:2_ determined via the traditional GC analyses were mostly higher than their pyrolysis counterparts. This may be due to the improved chromatographic separation between C_18:1_ and C_18:2_ FAMEs achieved by the longer 120 m column employed for traditional GC-IRMS analyses and/or the greater number of replicates analyzed via this protocol. α_W_ values calculated from the Pyr-GC-IRMS for ergosterol are inconsistent with their GC-IRMS counterparts, and might be due the lower signal for ergosterol detected via Pyr-GC-IRMS (Fig. 7) compared to the single, typically large peak observed with traditional GC-IRMS (not shown). Overall, our findings suggest that the Pyr-GC-IRMS protocol for determining stable C and H isotopic composition of C_18:2_ represents a robust and fast alternative to the conventional GC-IRMS analysis.

**Figure 7:**
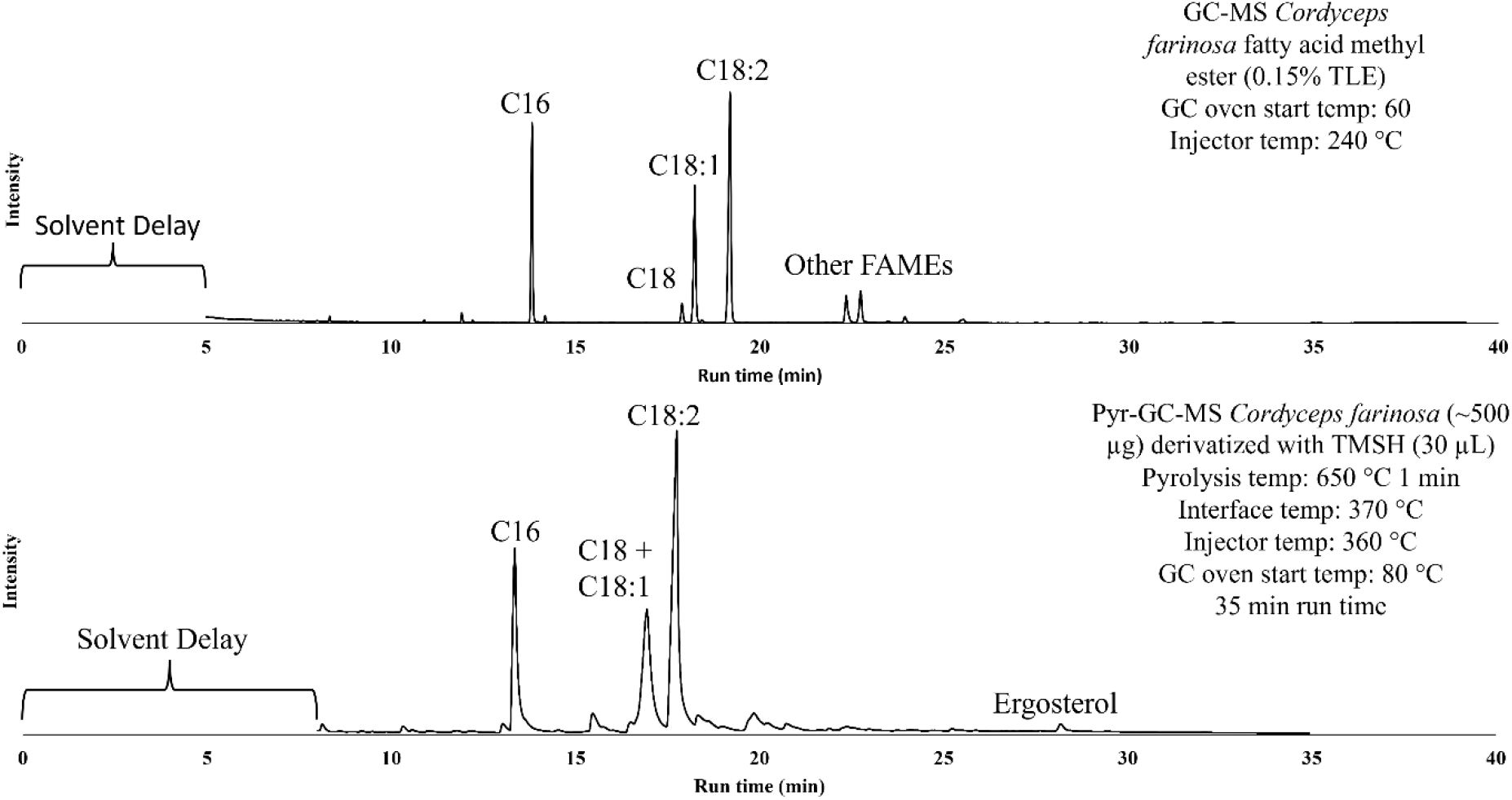
Typical FID chromatogram of a *Cordyceps farinosa* FAMEs/Lipids measurement with conventional GC-MS (top) and Pyr-GC-MS (bottom). Separation of FAMEs for the fungal biomarker C_18:2_ was successfully achieved with both methods. Advantage of the Pyr-GC method includes the measurement of both FAMEs and ergosterol in the dry biomass, whereas for the conventional method, sterols and FAMEs analysis are pre-extracted and analyzed separately.

An advantage of the Pyr-GC-IRMS approach was the tandem measurement of fatty acids and ergosterol in the same analytical injection (Fig. 1) which reduces the time of analysis, but requires high pyrolysis and GC temperatures for optimal detection^49^. On the other hand, Pyr-GC analyses required the use of a shorter separation column (30 m) and thus poorer peak resolution, because the pyrolysis unit could not withstand the higher backpressure of the 120 m column used for optimal separation of FAMEs. Nevertheless, the IL60 column achieved decent baseline resolution for C_18:2_ and allowed for detection of ergosterol in the same injection and negligible interference of other compounds in fungal biomass samples, owing to the high maximum operation temperature for this column (300 °C), which is a limitation for most polar columns (max temperature around 260 – 280 °C).

## 4. Conclusion and future perspective

The purpose of this work was to apply the dual SIP approach for determining heterotrophic CO_2_ fixation and water H assimilation into fungal lipid biomarkers when grown on labile substrates (glucose and glutamic acid) available in soils systems. Our findings suggest that fungi solicit distinguishable isotope effects during the incorporation of ^13^C-DIC and HDO into their membrane lipids when growing on glucose versus glutamic acid as the C substrate. We could show that the heterotopic IC incorporation into fungal biomarkers was within the expected range of 2 – 8%^35^ and we report for the first time the water H assimilation factor values for fungal biomarkers, which are consistent with α_W_ values reported for bacteria. Ergosterol was less informative as a fungal biomarker in these studies as in comparison to the more distinguishable signals exhibited by C_18:2_; however, it may serve as a potential marker for a distinction between labile and stable carbon sources. Future application of this dual SIP assay to explore fungal activity in environmental settings has the advantages that both isotope labels can be added to environmental incubations with minimal alteration of the natural habitat, fungal lipid production can be estimated by HDO incorporation alone, and the ^13^IC/α_W_ ratio can inform which C substrates support their growth. The potential of this approach for environmental studies can be improved with additional growth experiments that more comprehensively define ^13^IC/α_W_ ratios imparted during growth on more stable forms of soil organic carbon (e.g., lignin) across a larger diversity of fungal taxa. This effort will be supported by the applicability of pyrolysis coupled GC-MS and IRMS for direct analysis of fungal biomass demonstrated in this study.

## 5. Methods

### 5.1 Cultivation & Harvesting

Pure cultures of microscopic fungi (S-Tab. 1), *Aspergillus dimorphicus* (AD) strain BCCO 20_2442, *Cordyceps farinosa* (CF) strain BCCO 20_1579, *Penicillium janczewskii* (PJ) strain BCCO 20_0265, *Acremonium polycromum* (AP) strain BCCO 20_1588 and *Aspergillus brasiliensis* (AB) strain CCM 8222 and one yeast species *Yarrowia lipolytica* (YL) strain MUCL 054012 were incubated at 25 °C in the dark in 2 L Schott bottles containing 50 mL of a modified minimum media Czapek-Dox Broth^25^, which was inoculated (1 mL) with approximately 10^6^ spores per mL. The growth medium contained per liter: 4 g organic C source (C_6_H_12_O_6_, glucose and C_5_H_9_NO_4_, glutamic acid, tested separately), 3 g NaNO_3_, 1 g K_2_HPO_4_, 0.5 g MgSO_4_, 0.5 g KCl, 0.01 g FeSO_4_, and 0.09 g NaHCO_3_. The pH was adjusted to 7.3. Dual-SIP experiments were performed using ^13^C-bicarbonate (^13^C-DIC, NaH^13^CO_3_) and deuterated water (D_2_O) according to Tab. 1. Each fungal species was grown in triplicate with non-labeled substrates (treatment 1), with δ^2^H of the medium water adjusted to i) 100‰ δ^2^H and 10% of ^13^C-DIC (treatment 2), ii) 200‰ δ^2^H and 10% ^13^C -DIC (treatment 3), and iii) 400‰ δ^2^H and 10% ^13^C - DIC (treatment 4). The Schott bottles were kept closed to prevent the labeled ^13^C-DIC from outgassing. Headspace samples for CO_2_ analysis (2 mL) were collected every 2 – 3 days from each bottle into helium-flushed (10 minutes at 100 mL min^−1^) 12 mL exetainer vials (Exetainer, Labco Limited, UK), to monitor fungal growth and ensure O_2_ availability. For the experiments with the species AD and CF growing on glucose, 1 L bottles were used and neither CO_2_ concentration nor isotopic composition were measured for AD.

**Table 1:**
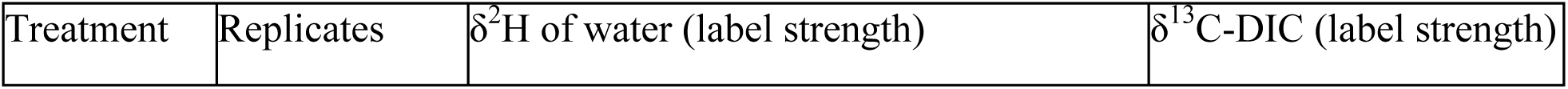
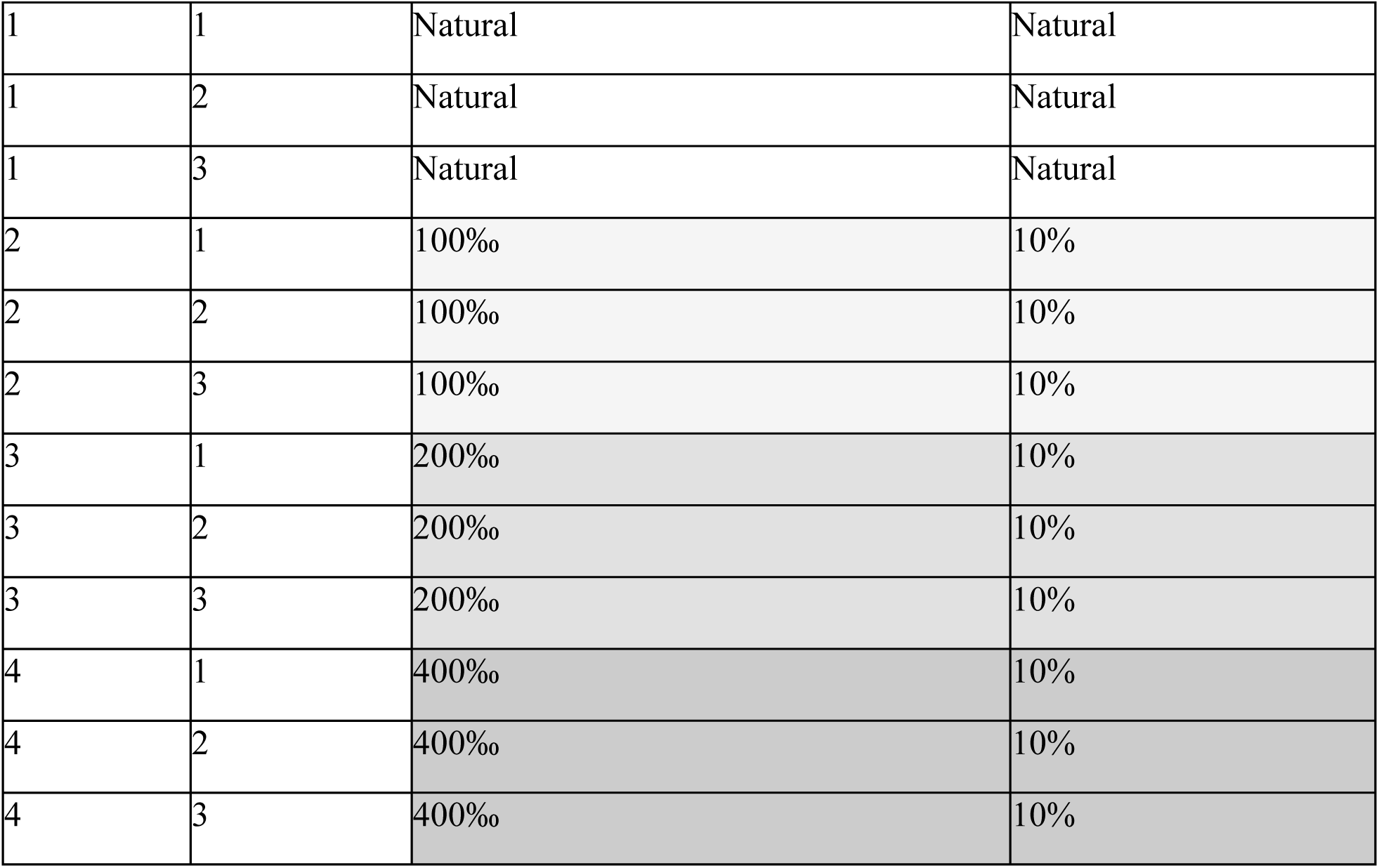
Incubation setup of fungal species for SIP experiments with ^13^C-bicarbonate (^13^C-DIC) and deuterated water (D_2_O).

| Treatment | Replicates | δ<sup>2</sup>H of water (label strength) | δ<sup>13</sup>C-DIC (label strength) |
| --- | --- | --- | --- |
| 1 | 1 | Natural | Natural |
| 1 | 2 | Natural | Natural |
| 1 | 3 | Natural | Natural |
| 2 | 1 | 100‰ | 10% |
| 2 | 2 | 100‰ | 10% |
| 2 | 3 | 100‰ | 10% |
| 3 | 1 | 200‰ | 10% |
| 3 | 2 | 200‰ | 10% |
| 3 | 3 | 200‰ | 10% |
| 4 | 1 | 400‰ | 10% |
| 4 | 2 | 400‰ | 10% |
| 4 | 3 | 400‰ | 10% |

At the end of the experiment the Mycelia were filtered from the growth medium using a sterile miracloth placed on a sterile funnel on a sterile 50 mL falcon tube. The mycelia on the miracloth were washed three times with sterile MilliQ water, then the mycelia were dried by first placing the miracloth on dry paper, then transferring to pre-weighed 50 mL falcon tubes. The biomass of the fresh (wet) weight was recorded and samples were stored at −20 °C until lyophilization, after which, the dry weight of each sample was determined and stored at −20 °C until further analysis.

#### 5.1.1 Growth monitoring

Headspace CO_2_ of fungal incubations (1 mL) were measured for CO_2_ concentration on a Hewlett Packard HP 5890 Series II GC (Agilent Technologies, Inc, Santa Clara, USA) coupled with a Thermal conductivity detector (TCD), equipped with a 30 m HP-PLOT/Q (polystyrene-divinylbenzene, 0.53 mm ID, 0.40 µm df, Agilent Technologies, Inc, Santa Clara, USA). The detector temperature was set to 130 °C, the oven was set to 70 °C and the system was operated with helium at a flow rate of 1 mL min^−1^.

### 5.2 Lipid extraction & chemical preparation

#### 5.2.1 Bligh & Dyer

Lyophilized fungal biomass was extracted using a modified Bligh and Dyer protocol^29^. Samples were sonicated for 10 min in four steps with a mixture of dichloromethane (DCM)/ methanol (MeOH)/ and an aqueous solution (1:2:0.8, v:v:v) by using 4 mL solvent per g sediment and extraction step. A phosphate buffer (8.7 g L^−1^ KH_2_PO_4_, pH 7.4) was used for the first two steps and a trichloroacetic acid buffer (50 g L^−1^ CCl_3_COOH, pH 2) for the final two steps. After each extraction step, the samples were centrifuged at 1250 rpm for 5 min and the supernatants were collected in a separation funnel. Phase separation was induced by addition of DCM and MilliQ and the organic phase was drawn off and collected in a Schott bottle. The aqueous phase was washed 3 more times with DCM, and then the pooled organic phase was washed 3 more times with de-ionized MilliQ water. Finally, the organic phase was collected as the total lipid extract (TLE) and was evaporated under a stream of argon and stored at −20 °C.

#### 5.2.2 Saponification

30% of the TLE was dried and saponified for fatty acid analysis, using 1 mL 6% KOH in MeOH for 3 h at 80 °C^30^. After cooling, 1 mL of 0.05 KCl was added. Neutral lipids were extracted four times using 1 mL n-hexane. The rest of the solution was treated with 10 % HCl to reach pH 1 and then free fatty acids were extracted four times using 1 mL n-hexane. Following that, all extracts were dried, mixed with 1 mL 20% BF_3_ in MeOH and heated at 70 °C for one hour to convert free fatty acids into fatty acid methyl esters (FAMEs). The solutions were extracted four times with 1 mL n-hexane. The combined extracts were evaporated under argon and stored at −20 °C until further analysis.

### 5.3 Measurements

#### 5.3.1 Deuterated water (HDO) isotope analysis

Liquid samples were transferred into 1.5 ml glass vials (32 x 11.6 mm, Fischer Scientific) and then measured using Triple Liquid Water Isotope Analyzer (Los Gatos Research), which is based on the principle of high-resolution laser absorption spectroscopy. Samples were dispensed into the instrument using an autosampler (PAL3 LSI, ABB company) and a 1.2 μL syringe (Hamilton). Samples were measured and evaluated against prepared laboratory standards of known isotopic composition. The isotopic ratios of these laboratory standards were verified by measuring against international standards (VSMOW2, SLAP2) made by the IAEA. To check the quality control, the measurements of the samples were also interspersed with periodic measurements of the prepared verification samples with known isotopic composition. The final isotopic composition (*δ*^2^H) was determined using LIMS software. Analytical error in case of *δ*^2^H was <1.5‰.

#### 5.3.2 Substrate analysis

Substrates (∼100 µg) were weighed into tin capsules (8 * 5 mm, Sercon, Crewe, UK) and placed in a helium-flushed carousel autosampler, then introduced to an Elemental Analyzer IsoLink IRMS system (EA IsoLink, Thermo Scientific, Bremen, Germany) equipped with a CHN/NC/N EA combustion/reduction reactor (Sercon, Crewe, UK) heated to 1020 °C. A pulse of oxygen was introduced to the reactor simultaneously with the sample. The sample gases were quantified via a thermal conductivity detector (TCD) and then introduced to a MAT 253 Plus isotope ratio mass spectrometer (IRMS; Thermo Scientific; Bremen, Germany) via the open split of a Conflo IV interface, with helium as the carrier gas. The isotopic composition was determined using Isodat 3.0 software against the corresponding CO_2_ working gas (−4.191‰ for δ^13^C), and the values were corrected for linearity and normalized to the VPDB scale using an international IAEA-600 (−27.771‰ for δ^13^C) standard. The analytical error was <0.04‰.

#### 5.3.3 Dissolved inorganic carbon (DIC) and CO_2_ isotopes

Medium water samples (2 mL) or CO_2_ headspace (2 mL) gas were injected into a helium-flushed 12 mL exetainer vials (Exetainer, Labco Limited, UK). Water samples were acidified with 0.1 mL 85% H_3_PO_4_ and left to equilibrate for at least 24 hours before measurement. Headspace CO_2_ was purged using a double-holed needle with helium into a 250 µL sample loop then separated via a Carboxen PLOT 1010 (0.53 mm ID; Supelco, Bellefonte, USA) held at 70 °C with a flow rate of 0.75 bar, and then introduced into the MAT253 Plus IRMS (Thermo Scientific, Bremen, Germany) via a Conflo IV interface. Each sample was injected three times during one analysis. The isotopic composition was determined using Isodat 3.0 software against the corresponding CO_2_ working gas (−4.191‰ for δ^13^C) and the values were corrected for linearity and normalized to the VPDB scale using internal house standards international standards IAEA-603 and IAEA-303A (2.46‰ and 93.3‰ for δ^13^C, respectively). The analytical error was <0.2‰.

#### 5.3.4 FAME analysis

FAMEs were measured using a GC Trace1310 gas chromatograph equipped with an SSL injector, a double gooseneck Topaz splittless liner (Restek, Bellefonte, USA) and a TR-FAME column (70% cyanopropyl polysilphenylene-siloxane; 120 m, 0.25 mm ID, 0,25 µm df, Thermo Scientific, Bremen, Germany). The samples were injected in splittless mode, with the injector temperature at 310 °C. The GC oven was held at 60 °C for 1 min, then ramped to 150 °C at 10 °C min^−1^, and to 250 °C at 4 °C min^−1^, then held at 250 °C for 15 min. Helium was used as carrier gas with a constant flow of 1.5 mL min^−1^. The column was coupled via a multi-channel device splitting the flow to a flame ionization detector (FID) and a mass spectrometer (MS), enabling both peak quantification and identification from one injection. The FID was set to 300 °C with flows of air at 350 mL min^−1^, H_2_ at 35 mL min^−1^, and N_2_ makeup gas at 40 mL min^−1^. The MS (ISQ QD; Thermo Scientific, Bremen, Germany) ion source was set to electron impact ionization mode (EI) at 70 eV and a scan range of m/z 50 – 500 with a scan time of 0.2 sec^−1^. The transfer line temperature was set to 280 °C and the ion source was set to 230 °C. The scan was started 15 min after the injection to avoid ionization of the solvent peak in the MS.

For FAME identification, the fragmentation patterns and mass spectra were compared to the National Institute of Standards and Technology (NIST) Spectral Library. Quantification was determined from the FID trace, according to the response of the internal standard (2-methyloctadecanoic acid; 2-MOA) and following Eq. 2.

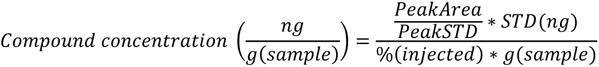

Equation 2: Final concentration is calculated with PeakArea being the integrated area of the unknown compound, PeakSTD the integrated area of the given standard and STD the known concentration of the standard in [ng], [%] the percent of the injected TLE and the mass of the extracted sample in [g]^29^.

Stable carbon and hydrogen isotope compositions of FAMEs were determined by splitting the flow from the GC column to a GC-Isolink II reactor, coupled to a MAT253 Plus IRMS via a Conflo IV interface; values are expressed in standard delta notation (δ^13^C and δ^2^H). MS information was simultaneously acquired by use of the multi-channel device described above. For conversion of FAMEs and ergosterol to CO_2_, the combustion reactor (nickel oxide tube with CuO, NiO, and Pt wires) was set to 1000 °C. For conversion of FAMEs and ergosterol to H_2_, the pyrolysis reactor (aluminum tube) was set to 1420 °C. FAMEs were identified by their retention time and fragmentation pattern. The isotopic composition was determined using Isodat 3.0 software against the corresponding CO_2_ or H_2_ working gas (−4.191‰ for δ^13^C, −239.5‰ for δ^2^H). Isotope corrections for instrument drifts, linearity, and normalization to the VPDB or VSMOW/SLAP scales were performed according to the response of USGS70 (−30.53‰ for δ^13^C, −183.9‰ for δ^2^H) and USGS72 (−1.54‰ for δ^13^C, 348.3‰ for δ^2^H) reference standards. The analytical error was <0.5‰ and <3.1‰ for δ^13^C and δ^2^H, respectively.

#### 5.3.5 Pyrolysis GC

The pyrolysis unit Shimadzu 3030D (Shimadzu, Kyoto, Japan/ Frontier Laboratories, Fukushima, Japan) was installed on top of the GC Trace1310 gas chromatograph SSL injector (Thermo Scientific, Bremen, Germany) and the GC was equipped with an SLB-IL60 column (non-bonded; 1,12-Di(tripropylphosphonium)dodecane bis(trifluoromethanesulfonyl)imide phase, 30 m, 0.25 mm ID, 0.20 µm df, Supelco, Bellefonte, USA). Due to the high restriction and thus backpressure from the 120 m FAME column, a different shorter analytical column (30 m) was used with the pyrolysis injector, which resulted in co-elution of C_18:0_ and C_18:1_ peaks (Fig. 1). The furnace temperature was to 650 °C and the interface temperature was set to 370 °C. The injector temperature was set to 360 °C and the GC oven was held at 80 °C for 1 min then ramped to 175 °C at 15 °C min^−1^, then ramped to 195 °C at 2 °C min^−1^ and then ramped to 300 °C at 10 °C min^−1^ and held for 7 min at 300 °C. Helium was used as carrier gas with a constant flow of 1.5 mL min^−1^ and split flow of 40 mL min^−1^. The column was coupled via a multi-channel device to enable simultaneous FID and MS acquisition from one injection. The GC-MS (ISQ QD; Thermo Scientific, Bremen, Germany) ion source was set to electron impact ionization mode (EI) at 70 eV and a scan range of m/z 50 – 500 with a scan time of 0.2 sec^−1^ was applied. MS scanning started after 8 min to avoid ionization of the solvent peak in the MS. Transfer line temperature was set to 300 °C and the ion source was set to 250 °C. The samples (lyophilized biomass, 80 µg – 1.5 mg) were weighed into an ultra clean stainless steel Eco-Cup LF (Frontier Laboratories, Fukushima, Japan), which were burned with a torch before usage to remove contaminants. FAMEs and ergosterol were detected in the same analytical run. Immediately prior to the measurement, 10 – 40 µL of trimethylsulfonium hydroxide (TMSH) was added on the sample to increase the volatization of the FAME and ergosterol and improve measurement sensitivity. Identification was performed using fragmentation patterns and the NIST 14 library. Quantification was performed with a calibration regression (0.5 – 40 ng) using an external standard (nonadecanoic acid, C_19_) for FAMEs and ergosterol (5 – 20 ng) for ergosterol. The same setup and method were also used to acquire stable carbon and hydrogen isotope composition of FAMEs and ergosterol via a GC-Isolink II device, as explained above. The column flow was split via a multichannel device to acquire MS and isotopic data simultaneously. The analytical error was <0.6‰ and <3.3‰ for δ^13^C and δ^2^H, respectively.

## Funding Sources

This Project was funded by the Czech Science Foundation GACR nr. 20-223805 and supported by MEYS CZ grant LM2015075 Projects of Large Infrastructure for Research, Development and Innovations as well as the European Regional Development Fund-Project: research of key soil-water ecosystem interactions at the SoWa Research Infrastructure (No. CZ.02.1.01/0.0/0.0/16_013/0001782).

## Acknowledgments

We thank Ljubov Polaková for the support of stable isotope measurements and laboratory protocols. We thank the Collection of Microscopic Fungi of the Institute of Soil Biology BC CAS (CMF-ISB) for providing the fungal species *Aspergillus dimorphicus* strain BCCO 20_2442, *Cordyceps farinosa* strain BCCO 20_1579, *Penicillium janczewskii* strain BCCO 20_0265, and *Acremonium polycromum* strain BCCO 20_1588, and the Czech Collection of Microorganisms (CCM) for kindly providing the *Aspergillus brasiliensis* strain CCM 8222. The authors declare no conflict of interest in the performance and publication of this research.

